# (Micro)saccade-related potentials during face recognition: A study combining EEG, eye-tracking, and deconvolution modeling

**DOI:** 10.1101/2023.06.16.545272

**Authors:** Lisa Spiering, Olaf Dimigen

**Author notes:** Address correspondence to: Olaf Dimigen, University of Groningen, Department of Psychology, Grote Kruisstraat 2/1, 9712 TS Groningen, The Netherlands. Author note: We thank Bagherzadeh-Azbari and colleagues (2023) for sharing the datasets analyzed in the present study. Supporting data and code are found at https://osf.io/e2rtp.

## Abstract

Under natural viewing conditions, complex stimuli such as human faces are typically looked at several times in succession, implying that their recognition may unfold across multiple eye fixations. Although electrophysiological (EEG) experiments on face recognition typically prohibit eye movements, participants still execute frequent (micro)saccades on the face, each of which generates its own visuocortical response. This finding raises the question of whether the fixation-related potentials (FRPs) evoked by these tiny gaze shifts also contain psychologically valuable information about face processing. Here we investigated this question by co-recording EEG and eye movements in an experiment with emotional faces (happy, angry, neutral). Deconvolution modeling was used to separate the stimulus-ERPs to face onset from the FRPs generated by subsequent microsaccades-induced refixations on the face. As expected, stimulus-ERPs exhibited typical emotion effects, with a larger early posterior negativity (EPN) for happy/angry compared to neutral faces. Eye-tracking confirmed that participants made small saccades within the face in 98% of the trials. However, while each saccade produced a strong response over visual areas, this response was unaffected by the face’s emotional expression, both for the first and for subsequent (micro)saccades. This finding suggests that the face’s affective content is rapidly evaluated after stimulus onset, leading to only a short-lived sensory enhancement by arousing stimuli that does not repeat itself during immediate refixations. Methodologically, our work demonstrates how eye-tracking and deconvolution modeling can be used to extract several brain responses from each EEG trial, providing insights into neural processing at different latencies after stimulus onset.

---

Under natural conditions, humans actively seek out relevant visual information with several eye movements per second. For example, while looking at a face, we might direct our gaze towards the most informative facial features to evaluate another person’s current emotional state. More generally, it seems that complex visual objects such as faces are not always fully processed during a single glance, but rather across multiple subsequent fixations (Hsiao, 2008).

In contrast to natural viewing conditions, electrophysiological (EEG) experiments on face recognition typically require participants to maintain a prolonged fixation. However, abundant evidence suggests that even under the strictest of fixation instructions, oculomotor exploration behavior continues at a miniature scale during EEG experiments in the form of small and often involuntary (micro)saccades (Yuval-Greenberg et al., 2008; Dimigen et al., 2009). These saccades typically remain unnoticed by the experimenter since the small rotation of the eyeballs produces corneoretinal artifacts that remain below common detection thresholds in the electrooculogram (EOG). In addition to these artifacts, however, each of the small gaze shifts also generates considerable visuocortical activity in the EEG (Dimigen, Valsecchi, Sommer, & Kliegl, 2009; Gaarder, Krauskopf, Graf, Kropfl, & Armington, 1964), at least if the stimulus is of sufficient size and contrast (Armington, Gaarder, & Schick, 1967; Gaarder et al., 1964).

In the current work, we used EEG, eye-tracking, and linear deconvolution modeling to test whether it is possible to fully separate these microsaccadic brain potentials from the temporally overlapping potentials elicited by the earlier stimulus onset. If possible, this would help EEG researchers to control for potential confounds from microsaccades in their data. More interestingly, however, in a second step, we also tested whether the fixation-related brain potentials (FRPs)^1^ generated by each of these small gaze shifts can be used to “probe” the ongoing state of neural stimulus processing during the trial. To explore this possibility, we tested whether the microsaccadic FRPs measured during a standard face recognition experiment are still sensitive to the facial expressions shown by the face in the same way as traditional, stimulus-locked ERPs (Schindler & Bublatzky, 2020). If this were to be the case, overlap-corrected microsaccade-induced brain activity could serve as new type of neural marker for attentional, cognitive, or affective processes.

## “Pinging” neural states with (micro)saccadic brain activity?

As noted above, there is now much evidence that during common EEG paradigms, visual areas are frequently re-activated by microsaccades (Dimigen et al., 2009) or small exploratory saccades (Dimigen & Ehinger, 2021) on the stimulus^2^. The most prominent component of the resulting FRP waveform is the lambda response, a positive potential that peaks at around 90 ms after the end of the gaze shift. The lambda response shares many features with the P1 in visually-evoked potentials (VEPs; Kazai & Yagi, 2003; G. W. Thickbroom, W. Knezevic, W. M. Carroll, & F. L. Mastaglia, 1991), especially to VEPs induced by pattern movement (G. Thickbroom, W. Knezevic, W. Carroll, & F. Mastaglia, 1991). It is clear that the lambda response is primarily visual in nature, as also evident by its sensitivity to low-level stimulus features (such as luminance contrast, Gaarder et al., 1964) and its absence or attenuation in darkness (Billings, 1989; Fourment, Calvet, & Bancaud, 1976). Nevertheless, the fact that microsaccade-related brain potentials are present in the vast majority of trials in common EEG paradigms (Dimigen et al., 2009; Meyberg, Werkle-Bergner, Sommer, & Dimigen, 2015) raises the questions of whether these potentials can be treated not just as artifacts (Yuval-Greenberg, Tomer, Keren, Nelken, & Deouell, 2008), but as a potential source of information.

One possibility is that the brain potentials produced by each of these small refixations are confined to early stages of visuocortical processing and that they lack cognitively-modulated or “endogenous” ERP components. This may also be due to rapid adaptation to the refixated stimulus (e.g., categorical adaptation of FRPs; Auerbach-Asch, Bein, & Deouell, 2020). Another possibility, however, is that FRPs from small saccades on the stimulus still reflect ongoing processes of attention and cognition. In the latter case, these potentials – if statistically separable from the stimulus-ERP – might actually be useful to “probe” the neural system at different latencies after stimulus onset.

Task-irrelevant but contrast-rich probe stimuli have been successfully used in EEG research to assess the locus of covert attention (Hillyard & Anllo-Vento, 1998; Luck, Fan, & Hillyard, 1993) and to decode otherwise “silent” working memory representations (Wolff, Ding, Myers, & Stokes, 2015; Wolff, Jochim, Akyürek, & Stokes, 2017). Since microsaccades generate visual transients that can be as strong as those from passive visual stimulation (Dimigen et al., 2009) they might be similarly useful for probing or “pinging” (Wolff et al., 2017) ongoing neural states. Evidence in favor of this idea comes from Meyberg et al. (2015) who analyzed microsaccades during the cue-target interval of an attentional cueing task. They found that both the VEPs to externally-flashed probes and the brain potentials from microsaccades reflected the cued locus of covert attention. Using the latter, it was also possible to dissociate covert attention, as reflected in the hemispheric lateralization of microsaccadic potentials, from overt attention, as reflected in the direction of microsaccades (for converging results, see Liu, Nobre, & van Ede, 2022).

These results indicate that it may be feasible to use the omnipresent microsaccades to obtain neural markers beyond those contained in the stimulus-locked EEG. However, whereas the microsaccades in the attention experiments described above typically occurred a few seconds after the last stimulus presentation, we first have to deal with the problem of overlapping potentials.

## Deconvolution modeling can separate overlapping responses

A major challenge for analyzing co-registered EEG/eye-tracking data is that the neural responses to stimulus onset overlap with those to subsequent fixations on the stimulus. Without correction, the stimulus-ERP waveforms will therefore be distorted by FPRs and vice versa (Coco, Nuthmann, & Dimigen, 2020; Gert, Ehinger, Timm, Kietzmann, & König, 2022). In addition to this overlap problem in the time domain, microsaccades can also affect stimulus-related EEG analyses in the frequency domain, for example by resetting (Dimigen et al., 2009; Gao, Huber, & Sabel, 2018) and lateralizing (Liu, Nobre, & van Ede, 2023) ongoing alpha oscillations.

A promising approach to address these overlap problems is linear deconvolution modeling, also known as finite impulse response deconvolution (Dale & Buckner, 1997). In this framework, within a large regression model, each observed EEG sample is understood as the summation of overlapping responses from different experimental events (Dandekar, Privitera, Carney, & Klein, 2012; Devillez, Guyader, & Guérin-Dugué, 2015; Dimigen & Ehinger, 2021; Kristensen, Rivet, & Guérin-Dugué, 2017b; N. J. Smith & Kutas, 2015b). Because the exact temporal interval between subsequent experimental events (e.g., stimulus onsets and microsaccades) varies naturally from trial to trial, it is possible to statistically separate the potentials related to each type of event. Additionally, the model accounts for both linear and nonlinear (Ehinger & Dimigen, 2019) influences of continuous event properties on the EEG; for example, it can account for the influence of saccade size on the FRP waveform (Yagi, 1979). After solving the model, the resulting regression coefficients can be analyzed like conventionally-averaged ERPs (N. J. Smith & Kutas, 2015a).

In summary, the linear deconvolution framework is promising to separate activity from multiple events within the same trial. In the current study, we build up on previous work (e.g., Dimigen & Ehinger, 2021; Kristensen et al., 2017b) to test whether we can fully separate stimulus- and microsaccade-related brain signals and whether the latter can then be used as a new type of probe for attentional and affective processing.

## Emotional facial expressions modulate stimulus-locked ERPs

In the current work, we explored these questions by focusing on the processing of emotional facial expressions. Effects of a face’s emotional valence (e.g., angry vs. neutral expression) are well-established in the literature where at least two prominent ERP components have been linked to the early and late processing of facial emotions (Schacht & Sommer, 2009; Schindler & Bublatzky, 2020):

The first and early component is the early posterior negativity (EPN), a negative deflection largest over bilateral occipitotemporal electrodes that differentiates emotionally neutral stimuli from those with a positive or negative valence (Schacht & Sommer, 2009; Schupp, Flaisch, Stockburger, & Junghofer, 2006; Schupp, Junghöfer, Weike, & Hamm, 2004). The EPN typically begins around 150 ms after stimulus onset and reaches a maximum between 200–300 ms (Schupp et al., 2006), although it can last up to 600 ms post-stimulus (Rellecke, Sommer, & Schacht, 2012). The EPN is larger for emotionally arousing stimuli (such as faces with an angry or happy expression) and this is commonly assumed to reflect a reflex-like allocation of attention towards emotionally arousing stimuli leading to their enhanced sensory encoding. It is often assumed that this reflects an innate predisposition for emotional faces to capture processing resources (Junghöfer, Bradley, Elbert, & Lang, 2001; Schacht & Sommer, 2009). Consequently, the EPN appears automatically regardless of the task or depth of stimulus processing (Rellecke, Palazova, Sommer, & Schacht, 2011; Rellecke et al., 2012). Since the EPN has consistently been shown to be modulated by facial expressions (Schindler & Bublatzky, 2020), we considered this component to be a suitable proxy to address our more general question of whether attentional or affective modulations are still present in the neural response elicited by microsaccades on the face^3^.

A second component linked to facial emotion processing is the late positive potential (LPP), a centroparietal positivity that emerges at around 350-500 ms post-stimulus (Schacht & Sommer, 2009; Schupp et al., 2006) and is larger for emotional stimuli. LPP effects have been observed in FRPs collected during the free viewing of emotional scenes, at least if the task requires an explicit arousal or valence rating (Simola, Le Fevre, Torniainen, & Baccino, 2015; Simola, Torniainen, Moisala, Kivikangas, & Krause, 2013). Unlike the EPN, the LPP is believed to reflect higher-level, elaborative, and nonautomatic stages of the encoding of emotional stimuli. As such, it is often absent if the task is superficial or if emotion is task-irrelevant, as it was the case in the current study. We therefore did not anticipate LPP effects in the present data.

## The present work

To summarize, the first aim of our study was to test whether we can use (non)linear deconvolution modeling to cleanly separate stimulus-locked responses from overlapping responses by small saccades. These genuine but often “hidden” cortical responses pose inferential hazards, since they are not removed by ocular correction algorithms (like independent component analysis, ICA). If saccade rates or directions differ between conditions, these potentials can also distort effects in stimulus-ERPs (Dimigen et al., 2009). Separating stimulus-from saccade-related brain activity should also lead to an improved signal-to-noise ratio.

Our second aim was to investigate whether after overlap-correction, the microsaccades themselves could be exploited as a source of information. More specifically, we examined whether the FRPs elicited by facial refixations merely reflect low-level changes in retinal stimulation or also a (re)processing of the face’s affective contents. In the latter case, we might be able to (1) extract multiple useful neural responses from each trial and (2) track the time course of affective processing via microsaccades at different latencies.

To address both questions, we reanalyzed data from a previously published experiment in which emotional expressions were displayed by static and dynamic faces (Bagherzadeh-Azbari, Lion, Stephani, Dimigen, & Sommer, 2022). We chose this experiment because it included simultaneous eye-tracking/EEG recordings and because faces were presented for 2000 ms in half of the trials. These relatively long trials allowed us to compare the neural responses following the first microsaccade on the face to those following microsaccades later in the trial. We hypothesized that the first microsaccade on the face – which typically happens 200-250 ms after stimulus onset (Engbert & Kliegl, 2003) – would still be crucial for ongoing face processing (Hsiao, 2008) and might therefore show the same arousal-related sensory enhancements as stimulus-locked ERPs (i.e., a larger EPN to happy and angry faces). In contrast, we expected EPN effects to be weak or absent for saccades late in the trial.

## METHODS

### Participants

We analyzed the data of a face classification experiment previously reported in Bagherzadeh-Azbari et al. (2022). In the study, twenty university students (12 female; age range: 18 to 44 years, *M* = 24.40, *SD* = 6.03) participated in the experiment for course credit or monetary remuneration. According to the Edinburgh Handedness Inventory (German version; Oldfield, 1971), all but one participant were right-handed (mean handedness score = +91.40, *SD* = 24.57) and all participants self-reported normal or corrected-to-normal visual acuity. Before participating in the experiment, participants provided written informed consent as approved by the departmental ethics review board of the Department of Psychology at Humboldt-University.

### Stimuli

Images of faces of 36 individuals (18 female, 18 male), were selected from the Radboud Faces Database (Langner et al., 2010). Each stimulus showed a frontal-view color image of a Caucasian face. External facial features (e.g., neck and hairline) were removed by a standard oval aperture (see Figure 1). During the experiment, each individual’s face was presented in nine different versions. It was shown with three different emotional expressions (neutral, angry, and happy) and also with three different gaze directions (the face looked directly at the observer, had an averted gaze position looking to the left, or an averted gaze position looking to the right). At the viewing distance of 60 cm, each face subtended 9.41° vertically and 7.07° horizontally. Both the size and the screen location of the presented faces were carefully standardized such that the eyes of the face always appeared in the same screen position across different trials. Figure 1B shows three example stimuli. More details on the creation and standardization of the faces are provided in Bagherzadeh-Azbari et al. (2022).

**Figure 1.**
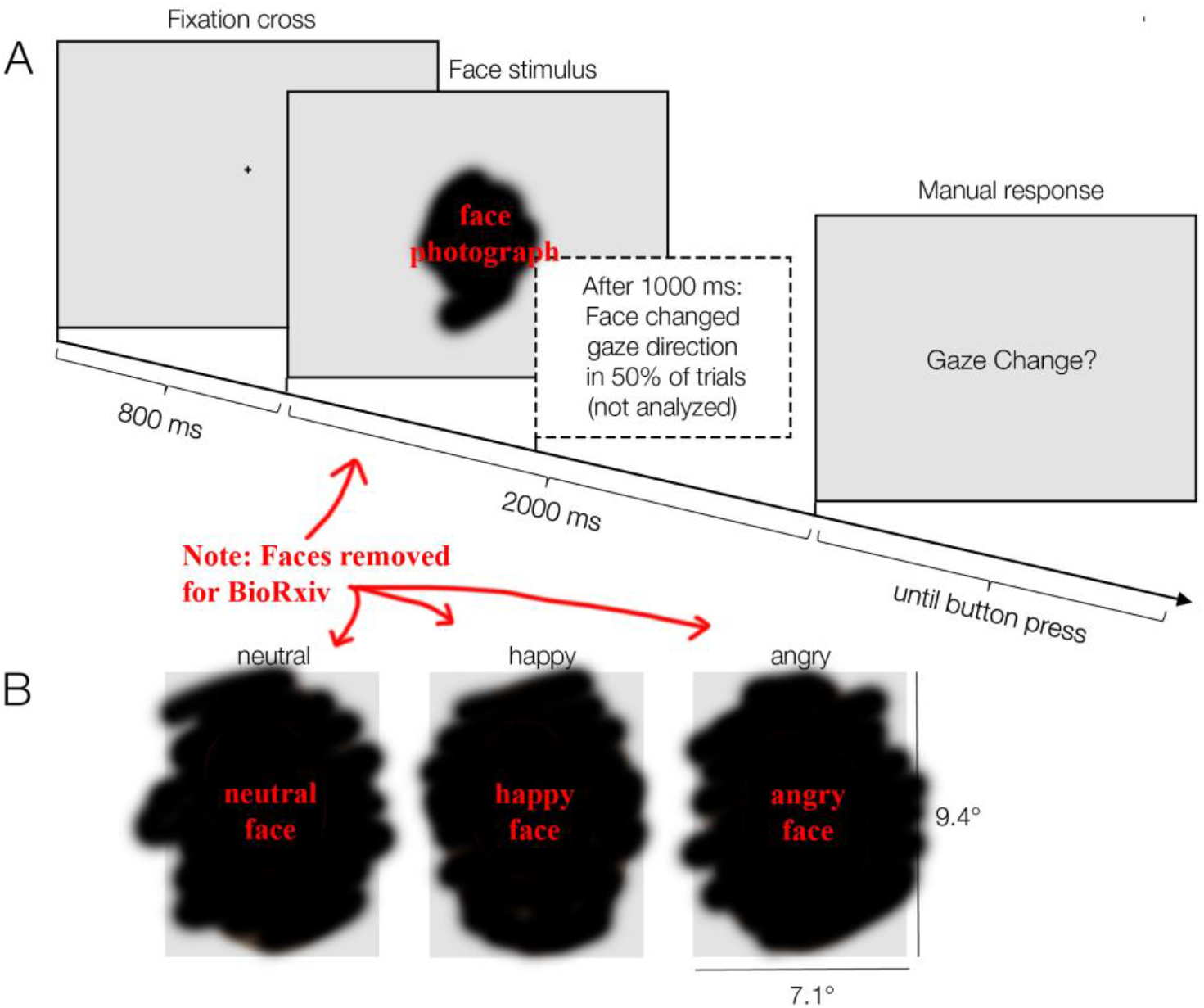
Trial scheme and example stimuli. **(A)** Participants were instructed to fixate a central cross shown for 800 ms. Afterwards, a face with a happy, angry, or neutral facial expression was presented for 2000 ms. The participant’s task was to indicate whether or not the gaze direction of the face (direct forward gaze, leftward, rightward) changed during the trial, which happened after 1000 ms in half of the trials (these trials were not analyzed here) **(B)** Stimuli varied with respect to two factors: emotional expression (happy, neutral, angry) and gaze direction (direct or averted gaze). Shown here are the three emotional expressions for a face with a direct (forward) gaze.

### Procedure

Participants were seated in an acoustically and electrically shielded cabin in front of a 22-inch cathode ray tube monitor (Iiyama Vision Master Pro 510, vertical refresh rate: 160 Hz, resolution: 1024 × 760 pixel). Following preparation of the EEG, participants first performed a 7-min calibration routine during which they made eye blinks as well as 15° eye movements in all four cardinal directions. These isolated saccades were later used by the ocular artifact correction procedure (see section *Preprocessing* below). This calibration task was followed by the face recognition experiment.

The trial scheme of this experiment is illustrated in Figure 1A. Each trial started with the display of a black fixation cross (0.72° × 0.72°) presented on a homogeneous gray background. The fixation cross was presented 1.44° above the screen center and therefore centered on the nasion (bridge of the nose) of the future face stimulus shown immediately afterwards; the initial viewing position on the face was therefore always in-between the eyes, near the optimal viewing locations for faces (Hsiao, 2008).

After 800 ms, the fixation cross was then replaced with the face stimulus. Half of the trials were static no-change trials, which were used for the current analysis. In these trials, the face was presented for 2000 ms and remained unchanged. The other half of trials were dynamic gaze-change trials, which were not analyzed here. In these trials, the face remained on screen for 1000 ms, but was then replaced for the remaining 1000 ms by an almost identical version of the face that only differed in terms of its gaze direction. For example, the face might initially look at the observer during the first second of the trial (direct gaze), but then avert the gaze to look away from the observer (or vice versa). In these gaze-change trials, only the eye region of the stimulus changed whereas the rest of the face – including the emotional expression – remained the same. After two seconds, the face was always replaced by a response screen (see Figure 1A), which prompted the participant to report whether the face had changed its gaze position or not.

As mentioned, for the purpose of the current study, we only analyzed the data from the no-gaze-change trials in which the face remained static for 2000 ms. Because of the long presentation duration of the faces in this condition, these trials provided an ideal opportunity to study the neural correlates of (micro)saccades executed at different latencies after stimulus onset.

The participant’s task was to watch the face and classify whether or not the face had changed its gaze direction during the trial. Participants were instructed to give their response only after the end of the trial using two manual response buttons operated with the left and right index fingers. The mean response accuracy was high (*M* = 97.8% correct, range across participants 92.8 to 100%, SD = 1.8%) and not further analyzed here. In case of incorrect or premature responses (before the stimulus face had disappeared), a red error message was shown. The participant’s button press initiated the next trial, which again started with the central fixation cross.

Participants received written instruction to focus on response accuracy, to fixate on the central fixation cross while it was visible, and to avoid blinking their eyes while the face was shown. Instead, they were encouraged to blink at the end of the 2-second face presentations.

The experiment comprised 864 trials, divided into 8 blocks, plus an additional 12 practice trials before the experiment. Facial emotion (neutral, happy, angry) and trial type (no-change vs. gaze-change) were counterbalanced and these six conditions were shown equiprobably during the experiment in a randomized order. Within the change-trials (not analyzed here), gaze changes leading to an averted gaze position (i.e., direct-to-averted gaze, left-averted to right-averted gaze, right-averted to left-averted gaze) and changes leading to a direct gaze position (i.e., left-to-direct, right-to-direct) occurred equally often. The same was true for averted gaze positions towards the left vs. right. The experiment was implemented using *Presentation*® software (version 18.10, Neurobehavioral Systems, Inc., Berkeley, CA).

### Eye movement recording

Binocular eye movements were recorded at a 500 Hz rate with a video-based eye-tracker (iView X Hi-Speed 1250, Sensomotoric Instruments GmbH, Germany). Head movements were restricted by the chin and forehead rest of the eye-tracker’s tower mount. The system was calibrated and validated with a 9-point grid before every block or whenever necessary during the experiment. Validations were accepted if the mean vertical and the mean horizontal validation error were both < 1°.

### Electrophysiological recordings

Electrooculogram (EOG) and EEG were recorded from 47 passive Ag/AgCl electrodes using BrainAmp amplifiers (Brain Products GmbH, Gilching, Germany). Forty-two electrodes were mounted on an elastic textile cap (EasyCap, Herrsching, Germany) at positions of the international 10/10 system; the exact montage is documented in the Online Supplement of Dimigen (2020). External electrodes were placed on the left (M1) and right (M2) mastoid; four EOG electrodes were placed at the outer canthus and infraorbital ridge of each eye. An electrode at FCz served as ground. Impedances were kept below 10 kΩ. To avoid pressure artifacts from contact with the eye-tracker’s forehead rest, foam rings were fitted around the prefrontal (Fp1/2) electrodes. During recording, all signals were referenced against electrode M1. Electrophysiological signals were sampled at 500 Hz at a time constant of 10 s with an online low-pass filter set to 100 Hz.

### EEG preprocessing and ocular artifact correction

Preprocessing of the EEG data was performed using EEGLAB (version 13.6.5b; Delorme & Makeig, 2004) and the EYE-EEG toolbox (version 0.81; Dimigen, Sommer, Hohlfeld, Jacobs, & Kliegl, 2011). In a first step, the EEG was digitally re-referenced to an average reference, thereby recovering the implicit reference (M1) as a recording electrode. Data was then bandpass-filtered between 0.1 to 45 Hz (−6 dB cutoffs) using EEGLAB’s windowed-sinc filter (*pop_eegfiltnew.m*) with default transition bandwidth settings. Ocular EEG artifacts were corrected using the Multiple Source Eye Correction method (MSEC; Berg & Scherg, 1994; Ille, Berg, & Scherg, 2002), as implemented in BESA (version 6.0; BESA GmbH, Gräfeling, Germany). The MSEC method provides an excellent correction of corneoretinal and blink artifacts (Dimigen, 2020) but only partially removes the saccadic spike potential, the sharp biphasic spike of synchronized extraocular muscle activity peaking at saccade onset (Keren, Yuval-Greenberg, & Deouell, 2010). Using the EYE-EEG toolbox, eye-tracking and EEG data were then synchronized based on shared trigger pulses sent frequently to both systems. The average synchronization error (misalignment of shared trigger pulses after synchronization) was < 1 ms.

### Trial selection

For data analysis, we focused exclusively on no-change trials during which a completely static emotional face was continuously shown for 2000 ms. This selection allowed us to study the FRPs elicited by saccades on the face at varying latencies without any confounds due to a change of the stimulus. Furthermore, in all analyses below, we aggregate our analysis across the gaze direction displayed by the face (direct, averted-left, or averted-right), since this factor was of no relevance for the current research questions.

In a first step, we identified clean trials in which neither the eye-tracking data nor the EEG contained missing data or artifacts from −200 to 2000 ms relative to stimulus onset. Three criteria were used to find clean trials: First, we rejected trials that included either an eye blink or gaze measurements outside of the stimulus image. Second, we excluded trials in which the mean gaze position during the 200 ms pre-stimulus interval was not within an invisible quadratic bounding box (side length: 3°) centered on the fixation cross (post-hoc fixation check). Finally, we discarded trials that contained remaining non-ocular EEG artifacts (after ocular artifact correction), defined as any voltages exceeding ±120 µV relative to the baseline voltages at each channel. In the remaining clean trials (80% of all analyzed trials), saccade and fixation events were detected using the binocular version of the velocity-based microsaccade detection algorithm by Engbert & Kliegl (2003) as implemented in the EYE-EEG toolbox (velocity threshold: 5 median-based SDs, minimum saccade duration: 8 ms, binocular overlap required).

### Eye movement analysis

We analyzed possible effects of facial emotion on five aspects of eye movement behavior: Saccade rate, saccade amplitude, and saccade direction (angle), as well as the vertical and horizontal fixation location within the face. Saccade rate was analyzed globally as a function of the emotional condition. For the four remaining dependent variables, measures were computed separately for the refixation following the first microsaccade on the face (called the “first fixation” in the following) and for all subsequent refixations on the face. For each type of saccade (first vs. subsequent) the four dependent measures were analyzed as a function of the emotional expression (neutral, happy, angry).

Since some of our eye movement measures have complex multimodal distributions (e.g., saccade angle is a circular predictor), we compared the distribution of each dependent variable across the three levels of *Emotion* using pairwise non-parametric Kolmogorov-Smirnoff (KS) tests (i.e., angry vs. happy, happy vs. neutral, and angry vs. neutral). Per dependent variable, we corrected the resulting *p*-values using the Bonferroni-correction (*p*_corr_) to account for the multiple pairwise comparisons. Saccade rate was corrected for 3 pairwise comparisons (*Emotion* levels); all other eye movement measures were corrected for 6 pairwise comparisons (2 *Saccade Types* × 3 *Emotions*). We computed the mean and standard deviations of saccade angles using the *CircStat* toolbox for MATLAB (Berens, 2009).

### Linear deconvolution modeling

Stimulus- and fixation-related brain responses were modeled and statistically separated using linear deconvolution modeling with additional nonlinear spline predictors for some variables (for reviews see N. J. Smith & Kutas, 2015a, 2015b) as implemented in the *unfold* toolbox for MATLAB (Ehinger & Dimigen, 2019). In the following, we will only provide a brief and informal summary of this analysis approach. For details, the reader is referred to recent tutorial papers explaining this approach in detail (Dimigen & Ehinger, 2021; N. J. Smith & Kutas, 2015b). Technical details on the *unfold* toolbox are found in Ehinger & Dimigen, 2019. After that, we will document how we set up the specific model for the present analysis.

Compared to traditional ERP averaging, linear deconvolution modeling has two crucial advantages for analyzing experiments with eye movements (Auerbach-Asch et al., 2020; Dandekar et al., 2012; Devillez et al., 2015; Dimigen & Ehinger, 2021; Gert et al., 2022; Guérin-Dugué et al., 2018). First, the normal temporal variability between different oculomotor events (e.g., stimulus onsets and saccade onsets) can be used to statistically disentangle the overlapping brain responses produced by each type of events. Second, the model allows the researcher to statistically control for the effects of various nuisance variables that are known to influence eye movement-related brain responses (such as the saccade amplitude preceding a fixation). Because these waveforms are estimated within a regression framework rather than with classic averaging, they are sometimes referred as regression-ERPs (rERPs, N. J. Smith & Kutas, 2015a), or, in the case of fixation-related potentials, as regression-FRPs. In the following, we will therefore refer to “rERPs” or “rFRPs” whenever we refer to waveforms obtained with the *unfold* toolbox.

One practical prerequisite of using linear deconvolution models is that the EEG recording is still continuous rather than cut into epochs. This is necessary so that the overlapping activity between all relevant experimental events can be considered in the estimation. To reduce computation time and the number of estimated parameters, the continuous artifact-corrected EEG was down-sampled to 200 Hz. In this continuous EEG dataset, we then included the stimulus onset events (coding the face onset event at the start of the trial) and the onsets of (re)fixations following microsaccades within the face. We only imported events from “clean” trials without missing data or residual artifacts (see above for screening criteria). To obtain the estimates for the non-overlapped rERPs and rFRPs, each channel of the continuous EEG signal was then modeled using a time-expanded design matrix, using the *unfold* toolbox. Time-expansion means that we model the continuous EEG in a certain time window (here: −200 to 800 ms) around each experimental event (here: stimulus onset and fixation onsets). For each time points, for each type of event (stimulus or fixation) and for each predictor in our regression model (e.g., the emotion of the viewed face), we then add a column to the design matrix which codes the state of this predictor at this time point relative to the event. This time-expansion step, illustrated in Figure 2 of Ehinger and Dimigen (2019), makes it possible to account for temporally overlapping effects of past and future events. This large regression model is then solved for the regression coefficients (or “betas”), which capture how much each event/predictor contributed to the measured EEG within the time expansion window. The resulting betas can be plotted like an ERP waveform (N. J. Smith & Kutas, 2015a).

**Figure 2.**
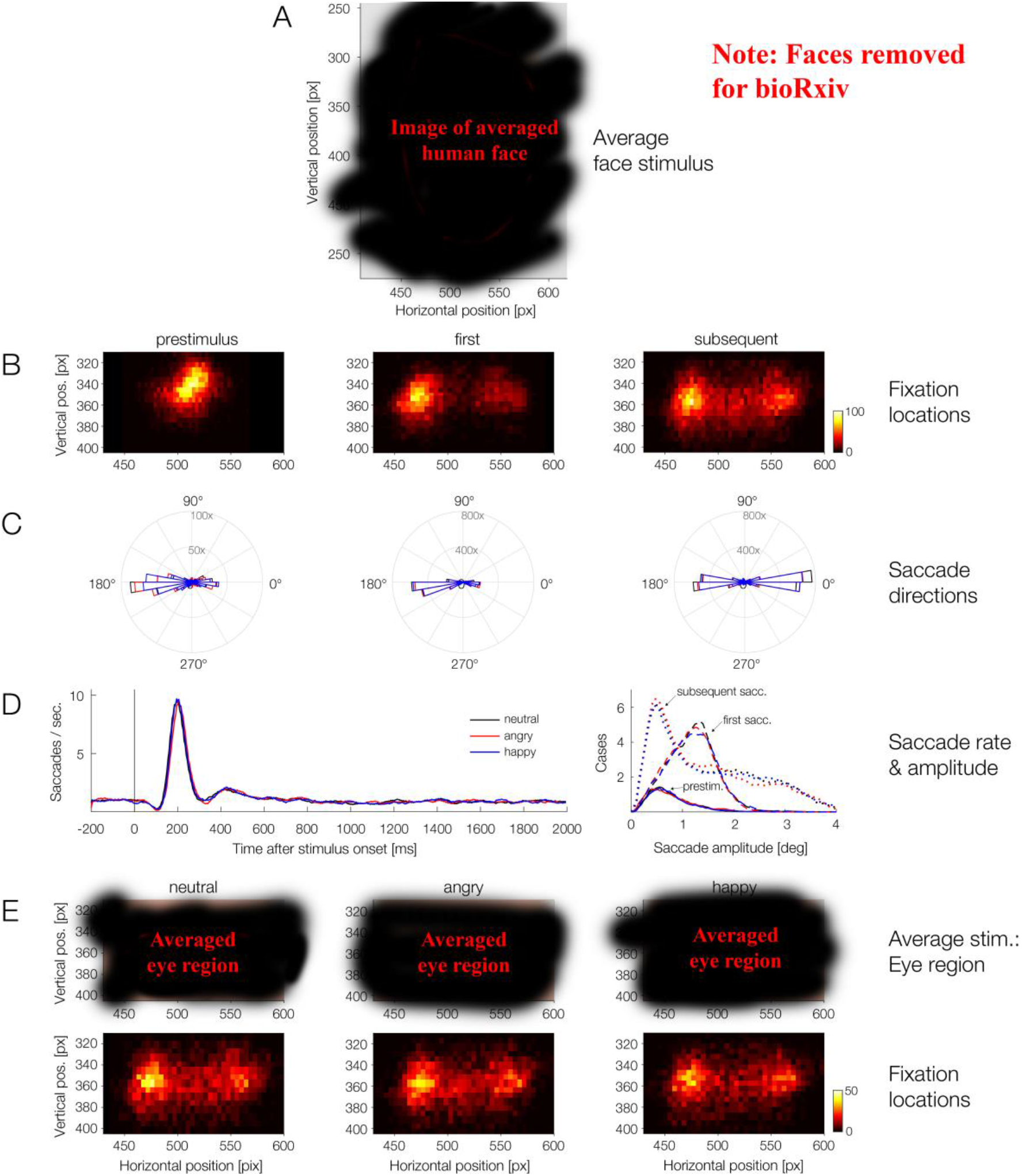
Properties of (micro)saccades detected during the trials. **(A)** Average of all presented face stimuli. The dashed rectangle around the eye region highlights the region for which fixation density heatmaps are shown in panels (B) and (E). Most saccades occurred within this eye region. **(B)** Fixation density plots (“heatmaps”) of fixation locations during the pre-stimulus interval (−200 to 0 ms, left column), for the first saccade (middle) and for all subsequent saccades during the trial (right). **(C)** Polar shots show the distribution of saccade directions for the pre-stimulus saccades, the first saccades, and for subsequent saccades. **(D)** Rate of saccades over time (left) and distribution of saccade amplitudes (right). Results are shown separately for the three emotion conditions. **(E)** Fixation heatmaps, shown separately for the neutral, angry, and happy condition across the post-stimulus interval (0-2000 ms).

Some predictors, such as saccade amplitude, have a strongly non-linear influences on the neural response in the EEG. Within the linear deconvolution framework, it is also possible to account for predictors that have nonlinear effects (N. J. Smith & Kutas, 2015b) by modeling their effects via a basis set of overlapping spline functions (cf. generalized additive modeling, GAM). For detailed tutorial reviews on how to model nonlinear effects within the deconvolution framework, the reader is referred to Dimigen and Ehinger (2021); Ehinger and Dimigen (2019); and N. J. Smith and Kutas (2015b).

In the following section, for clarity, we report the predictors used to model the brain responses elicited by stimulus onsets and fixation onset events as two separate sub-formulas. Please note, however, that these two formulas together specify one large regression model in which the regression coefficients (or “betas”) for all event types and predictors are estimated simultaneously. To model rERPs and rFRPs, we specified the following model, using a modified Wilkinson notation:

1. For stimulus onset events: `rERP ~ 1 + cat(Emotion)`
2. For fixation onset events: `rFRP ~ 1 + cat(Emotion) * cat(SaccadeType) + spl(SaccadeAmp,5) + FovealLum`

Here, the rERP waveform following stimulus onset events is simply modeled by an intercept term (`1)` and the categorical predictor *Emotion*, with the three levels of emotion categorically coded in the design matrix as 0 for neutral, 1 for angry, and 2 for happy. With the default treatment coding used by the *unfold* toolbox, the beta coefficients for the intercept term (`1`) simply correspond to the average rERP waveform elicited by the presentation of a neutral face. The resulting beta coefficients for the predictor *Emotion* then describe the additional effect produced by showing an angry or happy face, respectively. The betas of this predictor are therefore equivalent to a difference wave in a traditional ERP averaging analysis (e.g., with treatment coding, the betas for emotion level “2” are analogous to a difference wave “happy minus neutral”).

Within the same model, the brain responses following fixation onsets were modeled by a more complex formula: To model rFRPs, we again included an intercept term (`1`) as well as the categorical predictor *Emotion*, coded in the same way as for stimulus onsets. In addition, we added the categorical predictor *SaccadeType* that encoded whether the fixation followed either the first saccade (0) or a subsequent (1) saccade on the stimulus. In line with our hypothesis that emotion effects in FRPs might be stronger for the first fixation (i.e., following the fixation following the first saccade), we also allowed for an interaction between the *Emotion* and *SaccadeType*.

Finally, two continuous covariates were included in the model to control for low-level nuisance effects on the rFRP: saccade size (or amplitude, *SaccadeAmp*) and foveal image luminance (*FovealLum*). It is well-established that saccade amplitude has a strong influence on the shape of the post-saccadic brain response (e.g., Armington & Bloom, 1974; Yagi, 1979), with larger saccades followed by larger neural responses. Because this relationship is highly non-linear (Boylan & Doig, 1989; Dandekar et al., 2012; Dimigen & Ehinger, 2021; Kaunitz et al., 2014; Ries, Slayback, & Touryan, 2018) we included saccade amplitude as a nonlinear predictor modeled by a set of five spline functions (Dimigen & Ehinger, 2021; Ehinger & Dimigen, 2019).

Finally, as a second nuisance variable, we included the approximate luminance of the face stimulus at the currently inspected image location, which is also known to affect the post-saccadic neural response (Armington et al., 1967), with more luminant and more contrast-rich foveal stimuli generating larger lambda responses. For each fixation, foveal face luminance was calculated within a circular patch with a radius of 2° centered on the current fixation location. Approximate foveal luminance was estimated by taking the mean of the channel-weighted RGB values (using MATLAB function *rgb2gray.m*) of all image pixels within this region. Because luminance is well-known to have a logarithmic influence on P1 amplitude, this predictor was first log-transformed and mean-centered, and then added as a continuous predictor (*FovealLum*) to the model.

By solving the regression model for the betas, we obtained a time series of beta coefficients for each predictor in the design matrix. To obtain a waveform that corresponds to a traditional ERP curve (e.g., for angry faces), we can simply sum up the respective betas and also include the intercept term, which capture the overall waveshape of the ERP. For plotting rFRP waveforms, the two included nuisance predictors (saccade amplitude and foveal luminance) were evaluated at their respective mean values.

### Comparison to ERP averaging

Deconvolved potentials were compared to those obtained with traditional averaging. A useful feature of the *unfold* toolbox is that in addition to the overlap-corrected data, it can also compute the ERP/FRP averages that would result from the traditional averaging of the data, without controlling for overlapping potentials and covariates. To obtain these traditional averages, the continuous EEG was first cut into epochs of −200 to 800 ms around stimulus onsets and fixation onsets, respectively. In a second step, we generated the same design matrix of the model as for the deconvolution modeling, but without the time expansion step to control for overlapping potentials and without adding the continuous nuisance variables (saccade amplitude and foveal luminance) to the design matrix. This simple mass univariate regression model (N. J. Smith & Kutas, 2015a), which is equivalent to simple averaging, was then again solved for the betas.

### Second-level EEG statistics

#### Analyses of variance

Both traditional averaging and deconvolution modeling provide waveforms at the single-subject level, which can then be tested for statistical significance at the group level. All second-level statistical analyses were performed on the deconvolved potentials. As a first approach, we computed repeated-measures ANOVAs in an a-priori defined spatiotemporal regions-of-interest (ROIs) using the *ez* package for frequentist ANOVAs (Lawrence, 2016). As outlined in the *Introduction*, emotion effects are most reliably found on the EPN and LPP components, but LPP effects are often constrained to tasks that require an explicit emotion classification (Rellecke et al., 2012). Since facial emotion was not task-relevant, we expected valence effects mainly on the earlier EPN component. To capture the EPN, we used as a ROI the average voltage at six occipito-temporal electrodes (P7/P8, PO7/PO8, PO9/PO10) in the time window from 200-300 ms, based on prior studies on the EPN for faces (e.g. Schupp et al., 2006).

The average voltage in this spatiotemporal ROI was computed for both the deconvolved rERP and rFRP waveforms and submitted to repeated-measures ANOVAs as a second-level (group) analysis. In a first step, we ran a global repeated-measures ANOVA to test for emotion effects across the stimulus onset and fixation-related brain potentials. In this global ANOVA, we predicted average voltage in our ROI by the two within-subject factors *Emotion* (3 levels) and *Event Type* (2 levels: stimulus-locked/ERP, or refixation-locked/FRP), as well as the interaction between the two main effects. For this purpose, we averaged the rFRPs across *Saccade Type* for each emotion and participant.

Since this global ANOVA produced a strong interaction between Emotion and event type (see *Results*), we subsequently ran separate ANOVAs for each event type: Here, the ANOVA for stimulus-rERPs only included the three-level factor *Emotion*. The ANOVA for the fixation-rFRPs included the factors *Emotion* (3 levels), *Saccade Type* (2 levels: first or subsequent saccade) and the interaction term *Emotion* × *Saccade Type*. In case of significant effects, factor levels were compared with post-hoc *t*-tests.

#### Supplementary Bayesian analyses

As a supplementary analysis, we quantified the amount of evidence in favor or against emotion effects in rERPs and rFRPs. For this, we conducted Bayesian ANOVAs using the *BayesFactor* package for Bayesian ANOVAs in *R* (Morey, Rouder, Jamil, & Morey, 2015). An advantage of this Bayesian approach over a frequentist analysis is that it allows examining whether the data is more likely to have occurred under the null hypothesis (brain potentials do not differ between emotion conditions) or under the alternative hypothesis (brain potentials differ between emotion conditions).

The Bayesian ANOVA was calculated on the same EPN ROI window as the frequentist ANOVA. We first performed a factorial Bayesian ANOVA on stimulus-rERPs with one within-subject factor *Emotion*. For rFRPs, we first aggregated across *Saccade Type* and then ran a factorial Bayesian ANOVA with one within-subject factor *Emotion.* The resulting Bayes Factors (*BF*) were computed using the default Cauchy priors (*r* = 0.5 for the fixed effect of *Emotion*, and *r* = 1 for the random effect of subject). We used the classification by Lee and Wagenmakers (2013) to interpret the resulting *BF*s. The *BayesFactor* package estimates BFs with the Markov chain Monte Carlo (MCMC) algorithm (default number of samples = 10,000). Therefore, we report error percentages as an indication of numerical robustness of the BF. Here, lower error percentages reflect a greater stability of the BF, while error percentages below 20% are suggested as acceptable by van Doorn et al. (2021).

#### Cluster permutation tests

An additional cluster permutation test was used to also test for other emotion effects (Schindler & Bublatzky, 2020), outside of the predefined ROI and time window for the EPN component. Since still relatively little is known about emotion effects in FRPs (Guérin-Dugué et al., 2018; Simola et al., 2015; Simola et al., 2013), it is possible that their spatiotemporal properties differ from those in traditional ERPs (for a comparison, see Simola et al., 2013). For this purpose, we used the threshold-free cluster-enhancement (TFCE) procedure, a data-driven permutation test that stringently controls for multiple comparisons across time points and channels. The TFCE procedure was originally developed to address the multiple-comparison problem with fMRI (S. M. Smith & Nichols, 2009) but subsequently adopted to M/EEG data by Mensen & Khatami (2013). Compared to earlier variants of cluster permutation tests (Maris & Oostenveld, 2007) the main advantage of TFCE is that the researcher does not need to set an arbitrary cluster-forming threshold. Instead, the cluster-enhancement process of TFCE can be thought of as adopting all possible clustering thresholds.

For the present design, we used the ANOVA variant of the TFCE algorithm as implemented in the *ept_TFCE* toolbox (https://github.com/Mensen/ept_TFCE-matlab). The test was conducted across all 46 channels and the entire latency range from 0 to 500 ms after stimulus/fixation onset, respectively. We again applied this analysis only on the deconvolved potentials. Factors were specified in the same way as for the traditional repeated-measures ANOVAs described further above: The TFCE-ANOVA for rERPs only included the factor *Emotion*, whereas the one for rFRPs additionally included *SaccadeType* and the *Emotion* × *SaccadeType* interaction. Whenever the TFCE-ANOVA revealed significant effects or interactions of the three-level factor *Emotion*, we computed TFCE-based contrasts (*t*-tests) between the respective levels of the factor *Emotion*. Cluster-enhanced *F*- or *t*-values were compared against null distributions based on *n* = 5000 random permutations of the condition labels.

## RESULTS

### Eye movements

Simultaneous eye-tracking revealed that participants performed small saccades in the vast majority (*M* = 97.9%) of the trials. In trials with at least one saccade, the first saccade was usually followed (in 74.82% of all cases) by at least one more subsequent saccade on the face. Figure 2 summarizes the eye movements during the task. As shown in Figure 2, the majority of saccades was targeted at the eye region of the stimulus face. Figure 2B and 2C presents a more detailed visualization of the fixation locations at different latencies during the trial. During the pre-stimulus baseline interval (−200 to 0 ms), most saccades were located near the fixation cross, although participants already showed some tendency for making anticipatory saccades to the left (from the participants perspective) before stimulus onset. This is also visible in the polar histogram of saccade angles during the baseline period (see left polar plot in panel 2C; 64.87% of saccades were directed leftward).

Once the face stimulus was presented, the first saccade was almost always directed horizontally, with a clear preference for leftward rather than rightward saccades (see Nuthmann & Matthias, 2014 for similar results in natural scenes). As the center panel of Figure 2B shows, this initial saccade was typically aimed at the left eye of the face (69.76% of angles leftwards), and this left-eye bias was present in all three emotion conditions (see heatmaps in Figure 2E). Subsequent saccades on the faces, following the initial saccade, were also predominantly horizontally-oriented, but more widely distributed across the eye region, with a (slight) bias towards rightward saccades (52.69% directed rightwards, see Figure 2C right panel).

Saccades had a median amplitude of 1.23° (*SD* = 0.87), and this was quite similar for the first saccade (*M* = 1.21°, *SD* = 0.46) and subsequent saccades (*M* = 1.25°, *SD* = 1.02) on the face (Figure 2D right). The left panel of Figure 2D shows the rate of saccades over time. Across the entire duration of the trial, the mean rate was 1.29 saccades per second. Following the onset of the stimulus, we observed the typical biphasic pattern (Engbert & Kliegl, 2003) consisting of an initial suppression of saccades (saccadic inhibition), followed by a strong rebound: Almost no saccades were observed at around 120 ms after face onset (peak of inhibition) and this was followed by a strong increase in saccade rate, peaking shortly after 200 ms.

While the eye movements were generally highly similar for different facial emotions, a few eye movement parameters did show small but statistically significant differences between emotions. Table 1 shows the results of the KS-tests on all eye movement measures, analyzed across the post-stimulus interval from 0 to 2000 ms. In terms of the average vertical fixation location on the face, subsequent fixations on angry faces were located significantly lower on the face as compared to those on neutral or happy faces (angry-neutral: *D* = 0.05, *p_corr_* = 0.006; happy-neutral: *D* = 0.04, *p_corr_* = 0.03). However, these significant differences were again numerical extremely small (on average < 2 pixels or < 0.07° in both comparisons). Similarly, saccade amplitudes were slightly smaller in the angry as compared to both the neutral and happy conditions (first saccade: angry-neutral, *D* = 0.05, *p_corr_* = 0.02; subsequent saccade: angry-neutral, *D* = 0.04, *p_corr_* = 0.02; subsequent: angry-happy: *D* = 0.05, *p_corr_* = 0.001). Unstandardized effect size was again small; in terms of their median value, saccade amplitudes differed by less than 0.09° between emotion conditions. None of the other eye movement measures (saccade rate, saccade amplitude, and saccade angle, as well as horizontal or vertical fixation locations) showed significant effect of *Emotion* (all *p_corr_* > 0.05, see Table 1).

**Table 1.**
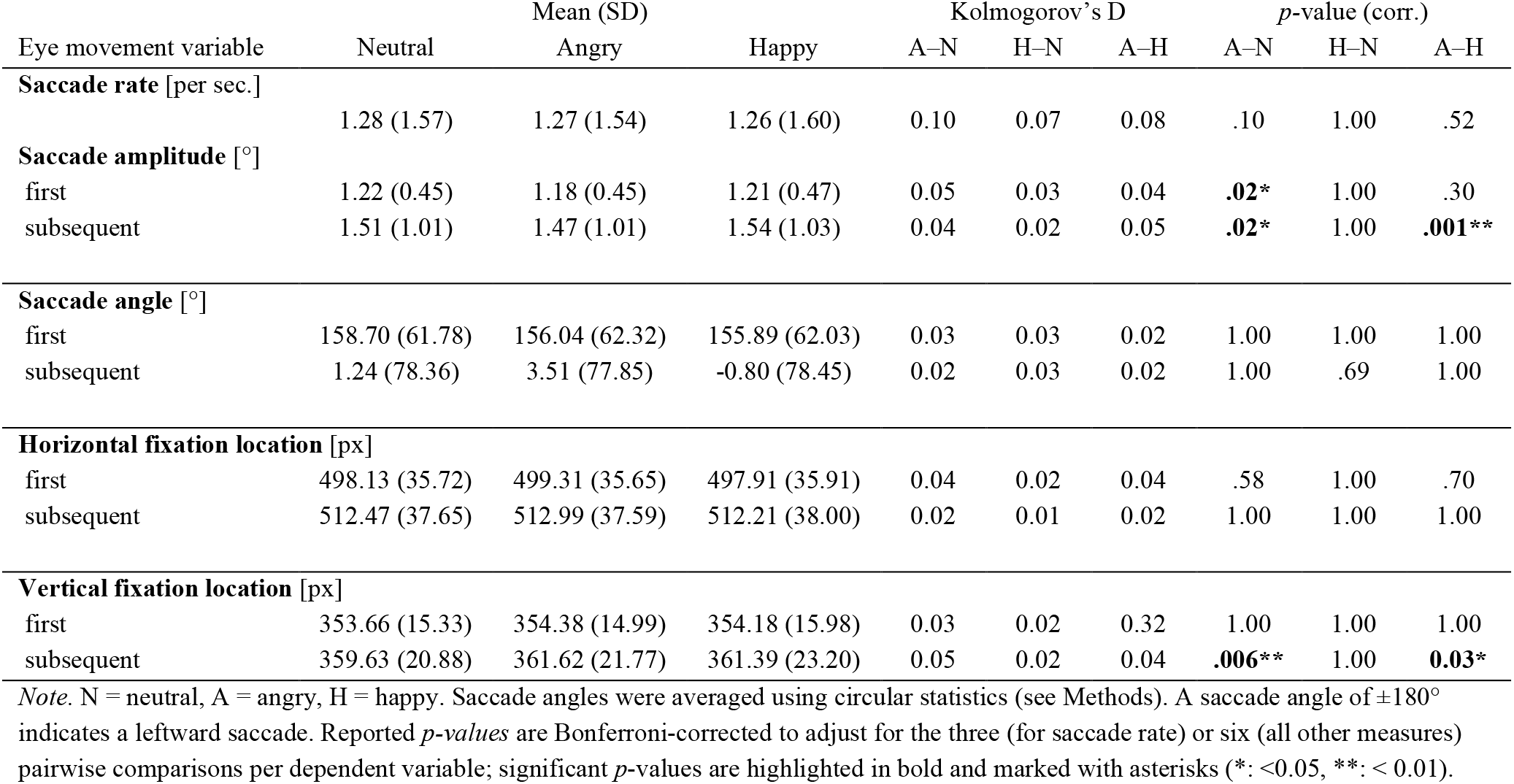
Test statistics for effects of *Emotion* on eye movements (KS-test results)

Taken together, the behavioral results show that in a traditional ERP experiment with a fixation instruction, participants made small saccades (of only around 1.2° amplitude) in virtually every trial. The majority of the oculomotor properties of these eye movements were not modulated by the emotional content of the faces; the unstandardized effect sizes of any significant emotion differences were small. Instead, participants’ eye movements were rather stereotypical, with the first saccade (after about 200 ms) being aimed at one of the eyes of the face, a highly task-relevant part of the stimulus. This first saccade was often followed by one or more subsequent saccades, which often stayed within the eye region of the face.

### Comparison of absolute brain responses with and without deconvolution

In most ERP studies, it is not considered that additional (micro)saccades happen during the trial. In a first step, we therefore assessed the impact of overlapping FRPs from these saccades on the waveshapes of the stimulus-locked ERPs and vice versa. In particular, we compared the waveforms obtained with traditional averaging (ERPs/FRPs) to those obtained with deconvolution modeling (rERPs/rFRPs).

#### Stimulus-ERPs

Compared to deconvolved rERPs, averaged ERPs show a distinct peak at around 300 ms (Figure 3A, upper panel), which was not present in the overlap-corrected data (Figure 3B, upper panel). With an amplitude of 2.81 µV, this distortion was most pronounced at electrode Oz and peaked about 314 ms post-stimulus (Figure 3C, upper panel). Since this occipital peak was eliminated by overlap-correction in the deconvolved rERPs, the measured distortion must originate from overlapping saccade-related potentials. This was confirmed by plotting latency-sorted single-trial stimulus-locked EEG epochs (*erpimages*). Without deconvolution (Figure 3A, lower panel), we can clearly see the confounding activity from the subsequent saccade on the face that is superimposed on the stimulus-induced ERP. This overlapping positivity, which corresponds to the eye movement-related visual lambda response, is successfully removed in the deconvolved EEG signals (Figure 3B, lower panel). Figure 3C shows only the “pure” overlapping fixation-related activity that had previously distorted the stimulus-ERPs. These results confirm previous reports on the contamination of stimulus-locked ERP waveforms by overlapping brain responses from (micro)saccades (Dimigen & Ehinger, 2021; Dimigen et al., 2009). Because the first saccade on the face occurs at a similar latency in most trials, the potentials do not jitter out, but create a considerable distortion of stimulus-ERPs at occipital scalp sites.

**Figure 3.**
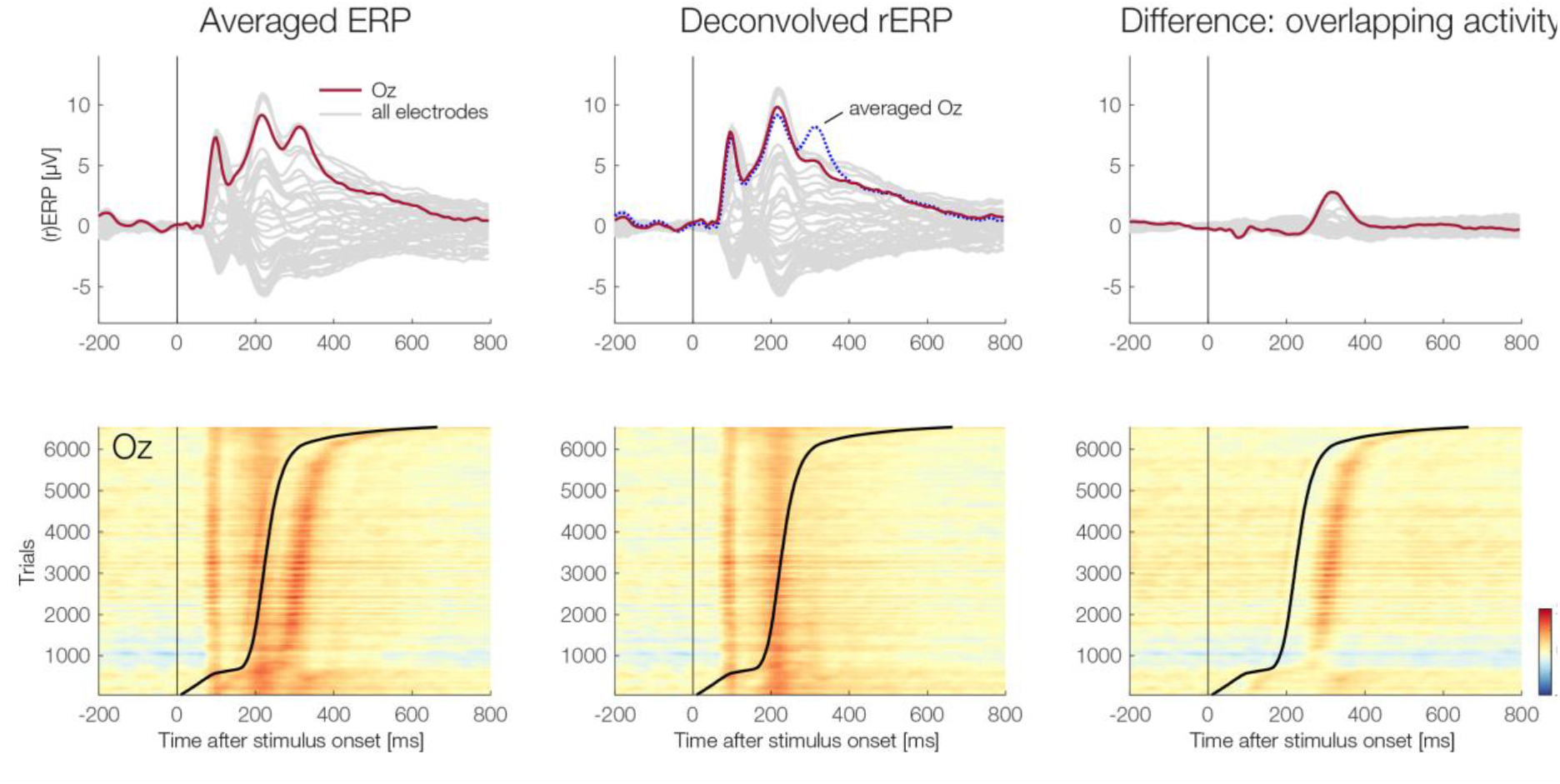
Comparison of stimulus-onset ERPs obtained with classic averaging (Averaged ERPs) and obtained with deconvolution (Deconvolved rERPs). **(A)** Averaged ERPs show a distortion at around 300 ms (upper panel). Single-trial latency-sorted EEG epochs at electrode Oz relative to stimulus onset (*erpimage*, lower panel). Trials were sorted by the latency of the first fixation onset on the stimulus (black sorting line). For clarity of visualization, single-trial epochs in *erpimages* were smoothed vertically across 100 adjacent epochs after sorting. **(B**) Deconvolved rERPs do not show the distinct peak at around 314 ms (upper panel). Latency-sorted single-trials after deconvolution suggest that the overlapping activity was successfully removed. Note that the latency-sorted trials of the deconvolved data include the model residuals, meaning that unmodeled overlapping activity would remain visible here if the overlap correction was incomplete. **(C)** The difference between the results without deconvolution (A) and with deconvolution (B). In the upper panel, the difference waves show the distortion introduced by overlapping eye movement-related brain activity. The lower panel shows the difference between the *erpimages* in panel A minus panel B and thus the “pure” overlapping activity elicited by the first eye movement on the stimulus (i.e., not modeled or unexplained EEG activity).

#### Fixation-FRPs

Compared to deconvolved rFRPs, averaged FRP curves were distorted before and after fixation onset (Figure 4A, upper panel). This distortion consisted of a smaller lambda response at 100 ms and drifting curves before and after. In contrast, deconvolved rFRPs show a clear spike potential and lambda response (Figure 4B, upper panel). The upper panel of Figure 4C shows the difference curves between averaging and deconvolution, highlighting the strong distortions in averaged FRPs. Deconvolved rFRPs varied according to preceding saccade type (first vs. subsequent) and according to saccade amplitude. Specifically, following the P1 (lambda response) peak, rFRPs elicited by the first saccade were more negative than those elicited by subsequent saccades, possibly reflecting adaptation (see Figure 4D). Our statistical analyses (reported in the next section) confirmed that this difference was significant, both in the EPN-ROI and the TFCE analysis. Saccade amplitude influenced the rFRPs in a nonlinear manner (Figure 4E): With increasing saccade size, the fixation-related P1 peaked earlier with a larger peak amplitude. However, as expected from previous research (Dandekar et al., 2012; Dimigen & Ehinger, 2021; Ries et al., 2018; Yagi, 1979), this increase with saccade amplitude was non-linear: A 0.5° increase in saccade amplitude lead to much larger change in P1 peak amplitude within the population of small (micro)saccades (i.e. from 0.5° to 1.0°) than within the population of more medium-sized saccades (i.e., from 1.5° vs. 2.0°). Figure 5 compares the brain potentials following stimulus-onset (rERPs) to those elicited by fixations (rFRPs). One interesting difference concern the N1 component: Whereas stimulus-evoked potentials showed a clear P1-N1 complex, the N1 was strongly attenuated or even absent in rFRPs (see arrows in Figure 5). More generally, rFRPs to refixations showed a striking absence of late/endogenous components in the waveform (Figure 5).

**Figure 4.**
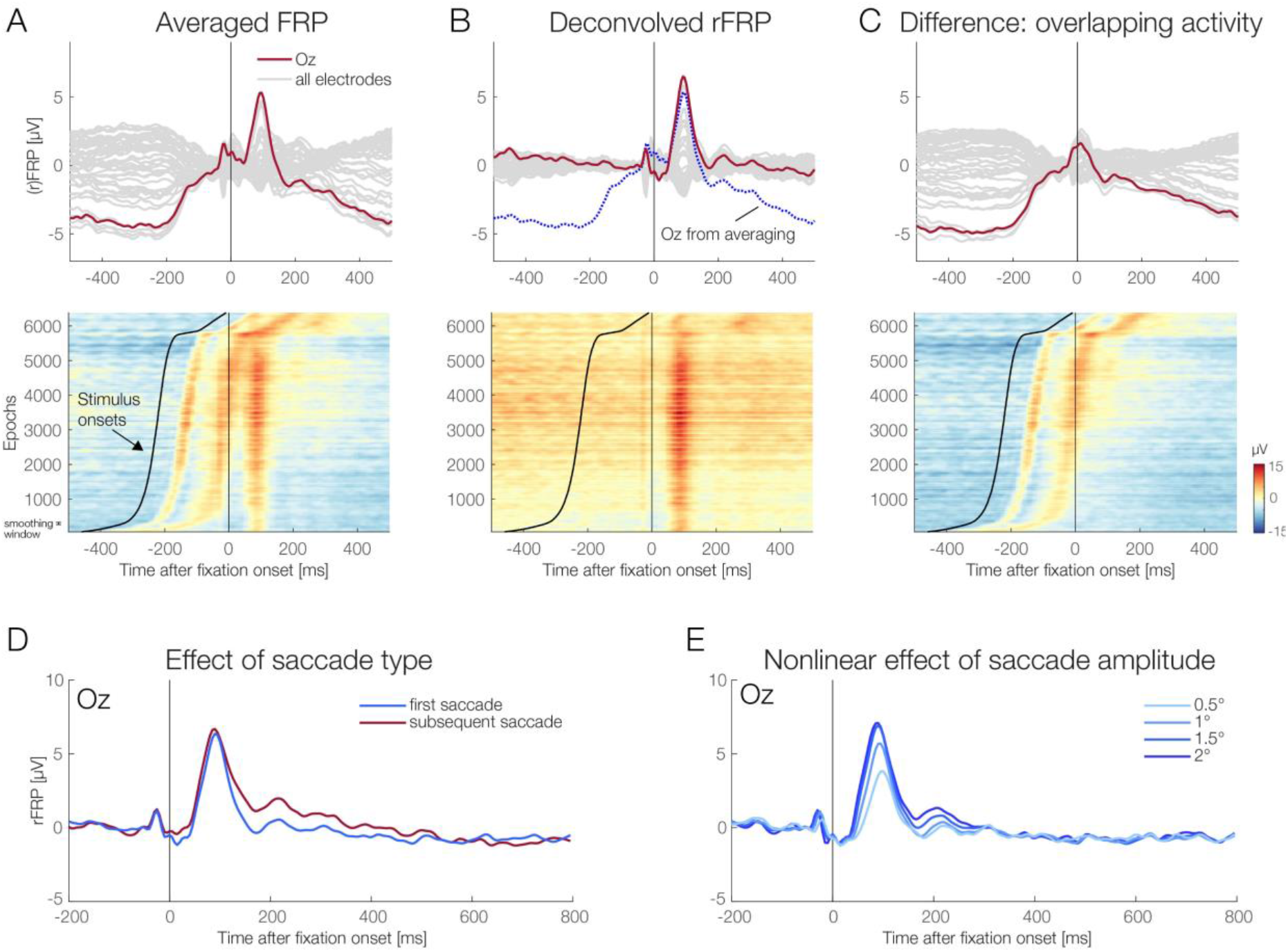
Comparison of fixation-related potentials obtained with classic averaging (FRPs) and with deconvolution (rFRPs). **(A)** Without overlap-correction, averaged FRPs are strongly distorted, both before and after fixation onset (upper panel). The lower panel shows the underlying single-trial EEG signals at Oz relative to fixation onset (*erpimage*). Trials are sorted by the latency of the preceding stimulus onset of the face on the screen, as indicated by the black sorting line. Data was smoothed vertically across 100 epochs after sorting. Strong distortions of the FRP waveform by the preceding stimulus onset are evident. **(B)** Overlap-corrected rFRPs do not show this distortion (upper panel). Lower panel show the corresponding *erpimage*, but now after deconvolution. This plot also includes the residuals of the model. **(C)** The difference between averaged vs. deconvolved FRPs shows the “pure” distortion of the FRP waveform by the preceding stimulus onset. **(D)** Effects of saccade type (first vs. subsequent) on rFRPs, illustrated at electrode Oz. Following the P1 peak, rFRPs for subsequent saccades remained more positive at occipital scalp sites. **(E)** Nonlinear effect of incoming saccade amplitude on rFRPs, illustrated at electrode Oz. Note that P1 amplitude is almost identical for 1.5° and for 2.0° saccades.

**Figure 5.**
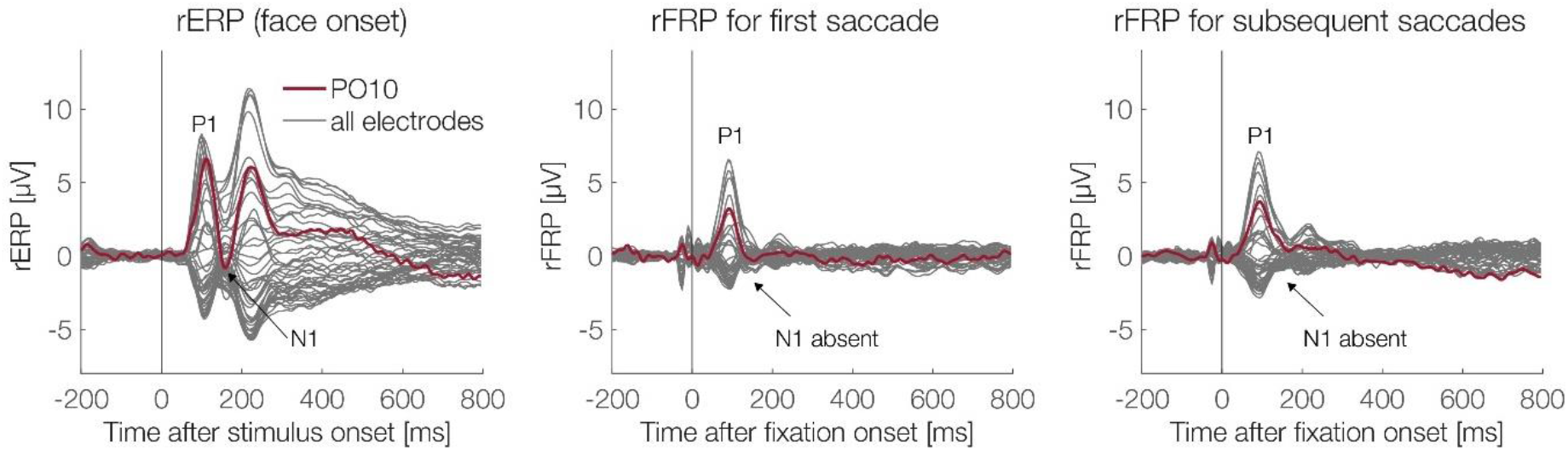
Lack of later components (e.g., N1/N170) in FRPs. The figure shows the grand-average overlap-corrected rERPs and rFRPs for the neutral condition at all EEG channels (dark grey lines). Electrode PO10, where N1/N170 effects to faces are often largest, is highlighted in red. Whereas the event-related potentials elicited by the onset of the face show a clear P1-N1 complex (left panel), the N1 is strongly attenuated or absent for the FRPs, both for the first fixation (middle panel) and for the subsequent fixations (right panel) on the face.

### Emotion effects in stimulus- and fixation-related potentials

In our study, we expected emotion effects on the EPN in both stimulus-ERPs and refixation-FRPs. Therefore, as a first step, we ran a repeated-measures ANOVA to test for emotion effects across even types, including both deconvolved rERPs and rFRPs. We applied this ANOVA on the average voltage in the predefined occipito-temporal ROI and in the a-priori defined EPN time window from 200-300 ms. For this global analysis, we also aggregated across the factor *Saccade Type* (first vs. subsequent saccades) in rFRPs. This global ANOVA revealed significant main effects of *Emotion*, *F*(2,38) = 3.87, *p* = 0.03, η_G_^2^ = 0.009, and *Event Type* (stimulus-onset vs fixation-onset), *F*(1,19) = 30.36, *p* < 0.001, η_G_^2^ = 0.45, as well as a significant interaction effect between *Event Type* and *Emotion*, *F*(2,38) = 5.95, *p* = 0.006, η_G_^2^ = 0.01. These results suggest that while there is a significant overall effect of emotion on neural responses in the EPN window, the effect also depends on whether these potentials are aligned to stimulus onsets or to refixations. In the next steps, we therefore analyzed the emotion effects separately within each event type.

#### Stimulus-related potentials (regression-ERPs)

Figure 5 depicts the deconvolved rERPs as a function on emotion condition. Consistent with our EPN hypothesis, the rERP waveforms, averaged across a region of interest for the EPN, were less positive for angry and happy versus neutral faces. The difference topographies of the three emotion conditions show the corresponding EPN-typical bilateral occipito-temporal negativity for the contrasts between the two emotionally-valence conditions (happy and angry) versus the neutral condition. We tested for emotion effects in rERPs in two ways: Firstly, we conducted ROI-based analyses based on our a-priori hypotheses on the spatiotemporal properties of the EPN component. Secondly, we used a TFCE-ANOVA that allowed us to find emotion effects outside our a-priori defined EPN time window and electrodes.

The classic ANOVA-based ROI analysis on the typical EPN time window (200-300 ms) revealed a significant main effect of *Emotion*, *F*(2,38) = 5.26, *p* = 0.009, η_G_^2^ = 0.03. Post-hoc frequentist *t*-tests revealed a significant difference between happy and neutral (*t*(19) = 2.82, *p* = 0.03, Cohen’s *d* = 0.63). Neither angry vs. neutral, *t*(19) = 2.61, *p* = 0.51, nor angry vs. happy, *t*(19) = −0.03, *p* = 1.00, differed significantly. Subsequently, we used a Bayesian ANOVA to quantify how much evidence there is for our hypothesized emotion effect in ERPs (and FRPs, see next section). We found moderate evidence towards the alternative hypothesis that there is an emotion effect (Bayes factor *BF =* 4.88, ±0.68%). An error percentage of 0.68% suggests strong robustness of the resulting Bayes factor. van Doorn et al. (2021) recommend percentages below 20% as acceptable, our percentages are way below this threshold.

The spatiotemporal properties of emotion effects in FRPs (Guérin-Dugué et al., 2018; Simola et al., 2015; Simola et al., 2013) are not yet as well-established as those in ERPs. Therefore, we used cluster permutation tests to also test for emotion effects across all electrodes and time points. The results of the TFCE-ANOVA on the differences among the within-subject factor *Emotion* on the rERPs are shown in Figure 5B. Convergent with the ROI-ANOVA, the main effect *Emotion* reached significance in the TFCE-ANOVA (peak: PO9 at 205 ms: *F*(2, 38) = 9.81, *p* = 0.03). Inspection of the TFCE plots (see Figure 5B) suggests that the overall effect in the TFCE is driven by a cluster of parieto-occipital electrodes (see Figure 5A) with smaller amplitudes for angry and happy compared to neutral starting beginning at around 160 ms and lasting for several hundred milliseconds. Post-hoc TFCE *t*-tests, visualized in Figure 5C, confirmed a significant difference between happy and neutral (peak significance observed at PO9 at 270 ms, *t*(19) = 3.67, *p* = 0.03, with amplitude difference of *M* = −1.38 µV) and between angry and neutral (peak significance at M1 at 175 ms, *t*(19) = 4.65, *p* = 0.049, amplitude difference of *M* = −1.20 µV). Angry and happy did not differ significantly in the TFCE analysis (*p* = 0.56).

Taken together, using both classic ROI- and TFCE-based ANOVAs, we found significant emotion effects in stimulus-locked regression-ERPs. Bayesian analyses suggest moderate evidence for emotion effects in rERPs. As expected, post-hoc frequentist ROI-based and TFCE *t*-tests revealed significantly lower amplitudes in happy vs neutral faces, while happy vs angry did not significantly differ. When comparing angry vs neutral faces, TFCE revealed a significant difference while ROI-based analyses did not. However, the *p-*value of the *t*-test from the ROI analysis was close to our pre-defined significance threshold (*p* = .051), and the TFCE results suggest that angry and neutral differed at time points and electrodes other than the ones we pre-defined in our spatiotemporal ROI.

**Figure 5.**
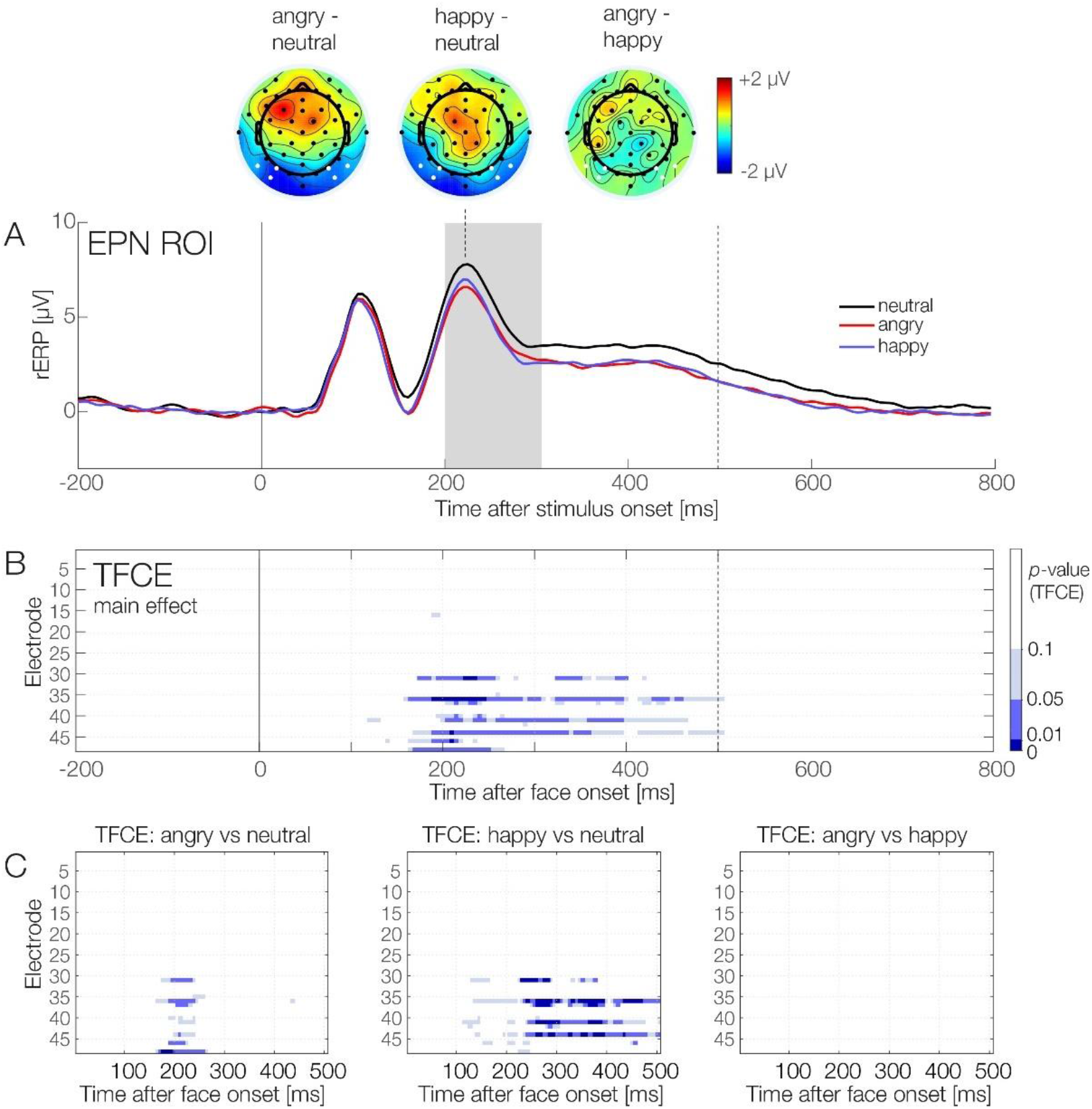
Emotion effects in deconvolved regression-ERPs. **(A)** Regression-ERP waveforms for the three emotion conditions, averaged over the a priori-defined ROI for the EPN component. Difference topographies contrasting the three emotion conditions are exemplified at the latency of 230 ms after face onset, electrodes belonging to the ROI are highlighted in white. The a priori-defined time window to quantify EPN amplitude (200-300 ms) is highlighted in grey. **(B)** and **(C)** Results of the cluster-permutation test (TFCE-ANOVA), conducted on the interval from 0 until 500 ms (dotted vertical line). Panel (B) shows the main effect of *Emotion*; the three panels in (C) show the post-hoc TFCE-based *t*-tests (lower panel) comparing the three facial expressions (angry, happy, neutral). The tests confirm the presence of an emotion effect, which distinguishes both the happy and the angry condition from the neutral condition. The contrast between the angry and happy condition was not significant.

#### Fixation-related potentials (regression-FRPs)

Figure 6 depicts the deconvolved rFRP curves as a function of emotion and saccade type. To test for emotion effects in rFRPs, we again used both a classic ROI-based and a TFCE approach. The classic ANOVA analysis in the predefined spatiotemporal ROI (occipito-temporal EPN ROI, 200-300 ms) revealed a significant main effect of *Saccade Type, F*(1,19) = 27.88, *p* < 0.001, η_G_^2^ = 0.06. This main effect of saccade type is also illustrated in Figure 4D, showing that the first refixation elicits more negative amplitudes at an occipital electrode (Oz) than subsequent saccades do. We found no main effect of *Emotion, F*(2,38) = 0.02, *p* = 0.98. The interaction between *Saccade Type* and *Emotion* was not significant either, *F*(2,38) = 0.26, *p* = 0.73. Using a Bayesian ANOVA in the same spatiotemporal ROI, we found moderate evidence against the hypothesis that there is an emotion effect in rFRPs (*BF =* 0.14, ±1.81%). Again, an error percentage of < 2% suggest strong robustness of this Bayes factor.

**Figure 6.**
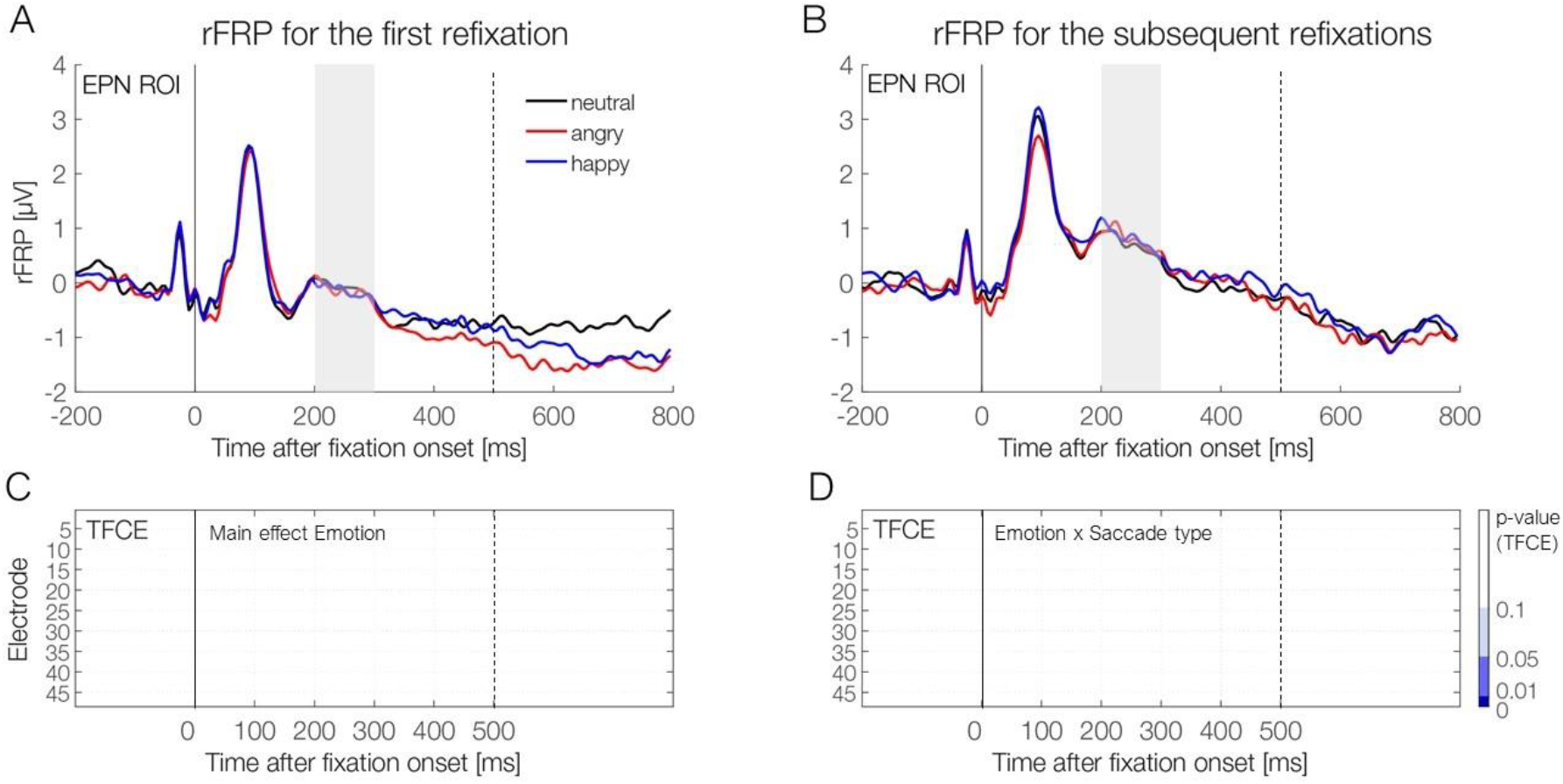
Regression-FRP waveforms for the three emotion conditions for **(A)** first refixation and **(B)** subsequent refixations on the face. Waveforms are shown averaged across the pre-specified ROI for the EPN component. The window used to quantify EPN amplitude (200-300 ms) is highlighted in grey. **(C-D)** Results of the cluster-permutation test (TFCE-ANOVA), conducted on the interval from 0-500 ms. (C) shows the non-significant main effect of Emotion, and (D) the non-significant interaction Emotion × Saccade type (first vs. subsequent saccade). These results suggest that there are no significant differences between the emotion conditions in rFRPs, regardless of whether the fixation followed the first or a subsequent (micro)saccade on the face.

The TFCE-ANOVA yielded convergent results as the classic frequentist ROI-based ANOVA analysis. The TFCE-ANOVA on rFRPs with *Emotion* and *Saccade Type* as within-subject factors revealed a significant main effect of *Saccade Type* (peak significance observed at electrode Iz at 160 ms; *F*(1,19) = 67.51, *p* < 0.05). However, there was no significant main effect of *Emotion* (*p* = 0.55), nor a significant interaction between *Emotion* and *Saccade Type* (*p* = 0.87). Taken together, while we find a classic emotion EPN effect in stimulus-evoked potentials, ROI and TFCE analyses provided no evidence that refixation-rFRPs are modulated by the emotional expression of a face.

## DISCUSSION

The active oculomotor exploration of the environment continues at a miniaturized scale and at a somewhat slower pace also during traditional EEG/ERP experiments. Previous work indicates that potentials from microsaccades may not just be a hidden source of artifacts (Yuval-Greenberg et al., 2008), but a source of useful information (Guérin-Dugué et al., 2018; Meyberg et al., 2015). In the current work, we applied linear deconvolution techniques to a traditional EEG face recognition experiment to separate brain potentials elicited by the stimulus onset from those generated by microsaccadic gaze shifts (i.e., small refixations) on the face. We hypothesized that each refixation would produce a volley of visuocortical activity (Dimigen et al., 2009) and we were interested whether an established ERP effect – that of emotional valence on the EPN component – would still be reflected in these FRPs. We hypothesized that this might be the case for the first fixation on the face, which usually occurs only around 200-250 ms after stimulus onset and therefore in the same latency range that the EPN emerges in stimulus-locked ERPs.

In our experiment, participants viewed emotional faces for two seconds with the task to report occasional gaze changes within the stimulus face. As expected, we found that rather small (median 1.23°) saccades on the face produced sizeable brain responses, whose amplitudes were at least similar to those elicited by stimulus onset, at least of the P1 component (cf. Figure 5). Importantly, linear deconvolution modeling allowed us to fully disentangle the stimulus-ERPs from these FRPs and vice versa. However, although we replicated the expected EPN effect of facial emotion in the (overlap-corrected) stimulus-ERPs, such an effect was not found in the FRPs elicited by microsaccades on the face. In the following, we will discuss our results in more detail, embed them into existing research, and provide an outlook for future research.

### Stereotypical saccades within the face were found in nearly every trial

Although a fixation cross was shown prior to face onset, participants made small saccades within the eye region of the face in virtually every (98%) trial. Following face onset, the saccade rate exhibited the common dynamic of an initial saccadic inhibition followed by a rebound (Engbert & Kliegl, 2003; Reingold & Stampe, 2002). Saccades rate reached a minimum at around 120 ms, but then rebounded strongly, with the first (micro)saccade typically happening after around 200 ms (Figure 2D).

Interestingly, the emotional expression shown by the face had only a minimal impact on these eye movements. While previous studies have shown that emotion can influence eye movements when participants are asked to categorize facial expressions (e.g., Eisenbarth & Alpers, 2011), our gaze shift monitoring task produced very limited differences in oculomotor behavior. Particularly, saccades on angry faces were slightly smaller and aimed slightly higher within the face (Table 1), but unstandardized effects sizes were small. Overall, the participant’s eye movement behavior was highly stereotypical: Beginning at the bridge of the nose, the first saccade was usually aimed at one of the eyes, usually the left one (from the perspective of the observer). While this gaze behavior was of course adaptive for the current change detection task, we observed similar stereotypical gaze behavior also in a previous experiment that required emotion classification (see Exp. 1 in Dimigen & Ehinger, 2021). We suspect that the repetitive presentation of hundreds of highly standardized face stimuli (no external features, identical screen location) explains this repetitive oculomotor behavior seen in EEG experiments on face recognition. Our finding that the size and direction of (micro)saccades was overall highly similar between emotion conditions should be reassuring for face researchers who are concerned about confounds from differences in gaze behavior between conditions (e.g., Vormbrock, Bruchmann, Menne, Straube, & Schindler, 2023).

As mentioned, participants exhibited a strong preference for looking at the left eye of the stimulus face, possibly due to a bias to extract information from the left visual field of another person’s face (Burt & Perrett, 1997; Gilbert & Bakan, 1973). Alternatively, this pattern may reflect a more general tendency to direct the first saccade on complex stimuli towards the left, a phenomenon which has been hypothesized to reflect a relative dominance of right-hemispheric parietal-frontal attention networks (“pseudoneglect”; Nuthmann & Matthias, 2014). Interestingly, we observed this bias already for microsaccades during the pre-stimulus baseline interval (see Figure 2C, left panel) showing that there is already an anticipatory attention shift towards the preferred left side.

In summary, we found that participants executed one or more small saccades towards the task-relevant eye region of the face in almost every trial. Most eye movements occurred stereotypically and synchronously at the start of the trial, with only marginal differences in oculomotor behavior between emotion conditions.

### Deconvolution cleanly isolates stimulus-from fixation-related potentials

Our first research aim was to use (non)linear deconvolution to separate ERPs from FRPs. To this end, we first compared traditionally-averaged ERPs with overlap-corrected regression-ERPs (rERPs). Over posterior scalp sites, averaged ERPs showed a large distortion of 2.81 µV from overlapping saccades peaking at 315 ms post-stimulus. The latency and amplitude of this distortion originated from the highly synchronous first saccade (at ~200 ms) which elicited a large lambda response about 90-100 ms later. A similar impact of microsaccades on P300 amplitude has previously been demonstrated (Dimigen et al., 2009). Importantly, after overlap correction, rERPs did not show this distortion anymore. Instead, the overlapping activity was cleanly separated, as evident by the absence of residual saccade-related activity in the latency-sorted single trials after deconvolution (see Figure 3B).

Secondly, we examined whether deconvolution can successfully isolate an rFRP waveform with a clear P1 (lambda response) and N1 component. As expected, without deconvolution, FRP waveforms were massively distorted by overlap (Coco et al., 2020; Gert et al., 2022). In contrast, deconvolved rFRPs showed none of these distortions anymore and also a clean and rather flat baseline (Figure 4D). We conclude that (non)linear deconvolution can successfully separate saccade-locked activity from stimulus-locked activity (see also Devillez et al., 2015; Dimigen & Ehinger, 2021; Gert et al., 2022; Kristensen et al., 2017b).

Besides the factor *Emotion*, we also included saccade amplitude as a nonlinear spline predictor in the model. Our results confirm a previously reported nonlinear relationship between saccade size and FRP amplitude (Dandekar et al., 2012; Dimigen & Ehinger, 2021; Dimigen et al., 2011; Dimigen et al., 2009; Ries et al., 2018; G. W. Thickbroom et al., 1991; Yagi, 1979). As shown previously (Dimigen & Ehinger, 2021), for unknown reasons, the influence of saccade size is highly nonlinear for the lambda response but more linear for later intervals of the rFRP waveform after about 150 ms (see Figure 4E). At a methodological level, these findings underline the importance of including saccade size as a nonlinear predictor in the model (Dimigen & Ehinger, 2021).

### Face onset-ERPs are enhanced by emotion, but refixations only reflect lower-level visual processing

Our second research question was whether both stimulus and fixation-related potentials would show an EPN effect of emotion. More specifically, we wanted to examine whether FRPs are enhanced by the reflex-like allocation of additional processing resources believed to underlie the EPN effect for arousing stimuli. In potentials time-locked to face onset, we observed the expected EPN effect from 200-300 ms, with more negative voltages at occipito-temporal electrodes for angry/happy faces as compare to neutral faces. Both the scalp distribution and timing of this effect resemble the EPN previously reported in the literature (Schindler & Bublatzky, 2020). The effect was also found in the cluster permutation test. In line with previous research (Rellecke et al., 2011; Rellecke et al., 2012), we found this EPN effect despite the fact that emotion was task-irrelevant. Although we did not formally test for emotion effects on the earlier N170, more negative voltages for angry and happy facial expressions were already observed seen during the peak of the N170 (see Figure 5A). It remains unclear whether this apparent effect is functionally distinct from the later EPN effect (Rellecke et al., 2013). There was no evidence that the later LPP component was modulated by emotion, which is expected since our experiment did not require an explicit emotion classification.

The core question was now whether a similar EPN effect – or any other effect of facial emotion – would also be seen in the brain responses elicited by refixations. This was not the case. The statistical analyses of rFRPs did not show a significant EPN effect of emotion, neither in rFRPs following the first saccade on the face, nor following subsequent saccades. The cluster permutation test, while less sensitive, also did not provide evidence for effects at other channels or time points. In support of this null result, a supplementary Bayesian analysis of the data found moderate evidence for the null hypothesis (no influence of emotion on rFRPs). In contrast, for the stimulus-locked ERPs, the Bayesian analysis provided moderate evidence in favor of the alternative hypothesis (emotion influences rFRPs).

In summary, our results indicate that once participants had been exposed to an emotional static face, they did not reprocess facial emotion when refixating it about 200 ms later. This apparently rapid processing of emotional facial expressions may seem surprising given that at least the recognition of faces seems to require more than one fixation (Hsiao, 2008). One likely interpretation of the current result is that the reflex-like allocation of more processing resources assumed to underlie the EPN effect only occurs once in response to the initial stimulus exposure. This is reminiscent of the rapid adaptation of the face-vs.-object N170 effect in FRPs (Auerbach-Asch et al., 2020) which is observed during the first but not during the second fixation of a (different) face stimulus during free viewing. Another possibility is that the potentials elicited by microsaccades are generally restricted to early cortical stages of the visual pathway. This interpretation would be consistent with the observation that the rFRPs in our studies show a strong lambda response (P1), but later components were seemingly attenuated or absent, including the N1/N170 component (see Figure 5).

Of course, whether or not microsaccadic potentials show attentional, cognitive, or affective modulations may strongly depend on the situation. In the current experiment, faces were static and facial emotion was both easy to process and task-irrelevant. In contrast to our results, Guérin-Dugué et al. (2018) observed an effect of emotional facial expressions on regression-FRPs during the free viewing of more complex and naturalistic face images (which also included the ears, hair, and some clothing). Specifically, these authors found significant differences only between surprised and neutral faces and only on the P1 (lambda response) and P2 components of the rFRPs elicited by the first refixation. There were no effects for the other emotion contrasts, e.g., those involving happy or disgusted faces. Surprisingly, these authors also did not observe any traditional stimulus-locked emotion effects (on the P1, N170, P2-P3 or LPP components) elicited by the face onset. One possibility to explain these discrepant findings might be the relative difficulty of extracting emotional cues from the faces in both studies. It is possible that the reflex-like sensory enhancements assumed to underlie the EPN only happen once during stimulus processing and that the timing of this enhancement depends on how difficult it is to decode the facial expression.

### Outlook

Our results show that existing techniques for EEG deconvolution modeling allow for a clean separation of stimulus-locked activity from the substantial but often unnoticed potentials generated by small gaze shifts on the stimulus. This not only makes it possible to eliminate potential confounds from stimulus-locked EEG measures, but also to extract multiple event-related brain responses from each trial of typical experiments. In the current work, we studied effects of emotion as a proxy to investigate the broader question of whether (micro)saccadic potentials contain psychologically meaningful information (Meyberg et al., 2015). Although we did not observe such an effect for the EPN component, it would be intriguing to examine similar effects in future research. For instance, it would be interesting to investigate whether other effects in mid-level vision, such as the N170 face inversion effect (Huber-Huber, Buonocore, Dimigen, Hickey, & Melcher, 2019) are also limited to the initial stimulus presentation or whether they recur for subsequent refixations. In other contexts, it may be possible to replace externally flashed probes with microsaccadic potentials, for example while probing the neural networks underlying working memory maintenance (Wolff et al., 2015). We hope that our study provides a useful framework for exploring (micro)saccade-related brain activity in diverse contexts in the future.

## OPEN PRACTICES STATEMENT

Supporting data and code are found at https://osf.io/e2rtp.

## Footnotes

1 Small saccades are very brief, so it is of relatively little consequence whether the EEG signal is aligned to the beginning or to the end of the movement. For the present work, we aligned the EEG signals to saccade offsets, that is, to the beginning of the (re)fixations produced by the saccades (fixation-related potential, FRP).

2 In the current work, we do not attempt to make a distinction between microsaccades and small exploratory saccades on the stimulus, but collectively refer to both as (micro)saccades.

3 Note that there is also evidence for topographically similar earlier effects of emotional expressions in the latency range of the N170 (Eimer & Holmes, 2007; Hinojosa, Mercado, & Carretié, 2015). However, some of these effects may not reflect a true modulation of the N170, but temporal overlap with the EPN (Rellecke, Sommer, & Schacht, 2013). While we focus on the EPN, we also tested for emotion effects across all latencies and channels.

## Notes

https://osf.io/e2rtp

### Competing Interest Statement

The authors have declared no competing interest.

